# Non-Negative Connectivity Causes Bow-Tie Architecture in Neural Circuits

**DOI:** 10.1101/2024.07.19.604347

**Authors:** Zhaofan Liu, CongCong Du, KongFatt Wong-Lin, Da-Hui Wang

## Abstract

Bow-tie or hourglass architecture is commonly found in biological neural networks. Recently, artificial neural networks with bow-tie architecture have been widely used in various machine-learning applications. However, it is unclear how bow-tie architecture in neural circuits can be formed. We address this by training multi-layer neural network models to perform classification tasks. We demonstrate that during network learning and structural changes, non-negative connections amplify error signals and quench neural activity particularly in the hidden layer, resulting in the emergence of the network’s bow-tie architecture. We further show that such architecture has low wiring cost, robust to network size, and generalizable to different discrimination tasks. Overall, our work suggests a possible mechanism for the emergence of bow-tie neural architecture and its functional advantages.

## Introduction

Bow-tie architecture (BTA) is ubiquitous in biological systems (*1–3*). BTA consists of substantially smaller or simpler intermediate systems linking between much larger and more complex systems. Hence, in terms of information processing, such an intermediate system can integrate large amounts of different information from different sources, generating a smaller amount of key information at the ‘waist’ of the architecture before re-using them for a wide variety of outputs.

BTA architecture can be found in various neural circuits, with smaller ensembles of neurons, for example within a brain region, receiving from and projecting to a larger number of neurons across wider brain circuits (**Fig. 1A-C**), resembling an hourglass structure (**Fig. 1D**). For instance, the mammalian retina is known to project to the primary visual cortex through the bottleneck lateral geniculate nucleus in the thalamic structure before projecting widely to cortical neurons, exhibiting a BTA architecture (*4–10*) (**Fig. 1A**). More generally, the thalamus has been known to be associated with ‘middleman-like’ information processing between larger brain regions, e.g. within the cortico-thalamic-cortical structure (*11–13*). Similarly, the olfactory system has a BTA network structure in which there is a large number of neurons with olfactory receptors in the antenna, projecting to a smaller number of neurons in the antennal lobe before the latter projecting to a widespread of neurons in the mushroom body (*14–18*) (**Fig. 1B**). Also exhibiting BTA are chemical neuromodulatory systems such as the relatively small midbrain dopaminergic nuclei, raphe serotonergic nucleui and norepinephrine locus coeruleus as compared to their wide afferent sources and efferent targets (*19–22*) (**Fig. 1C**).

**Figure 1:**
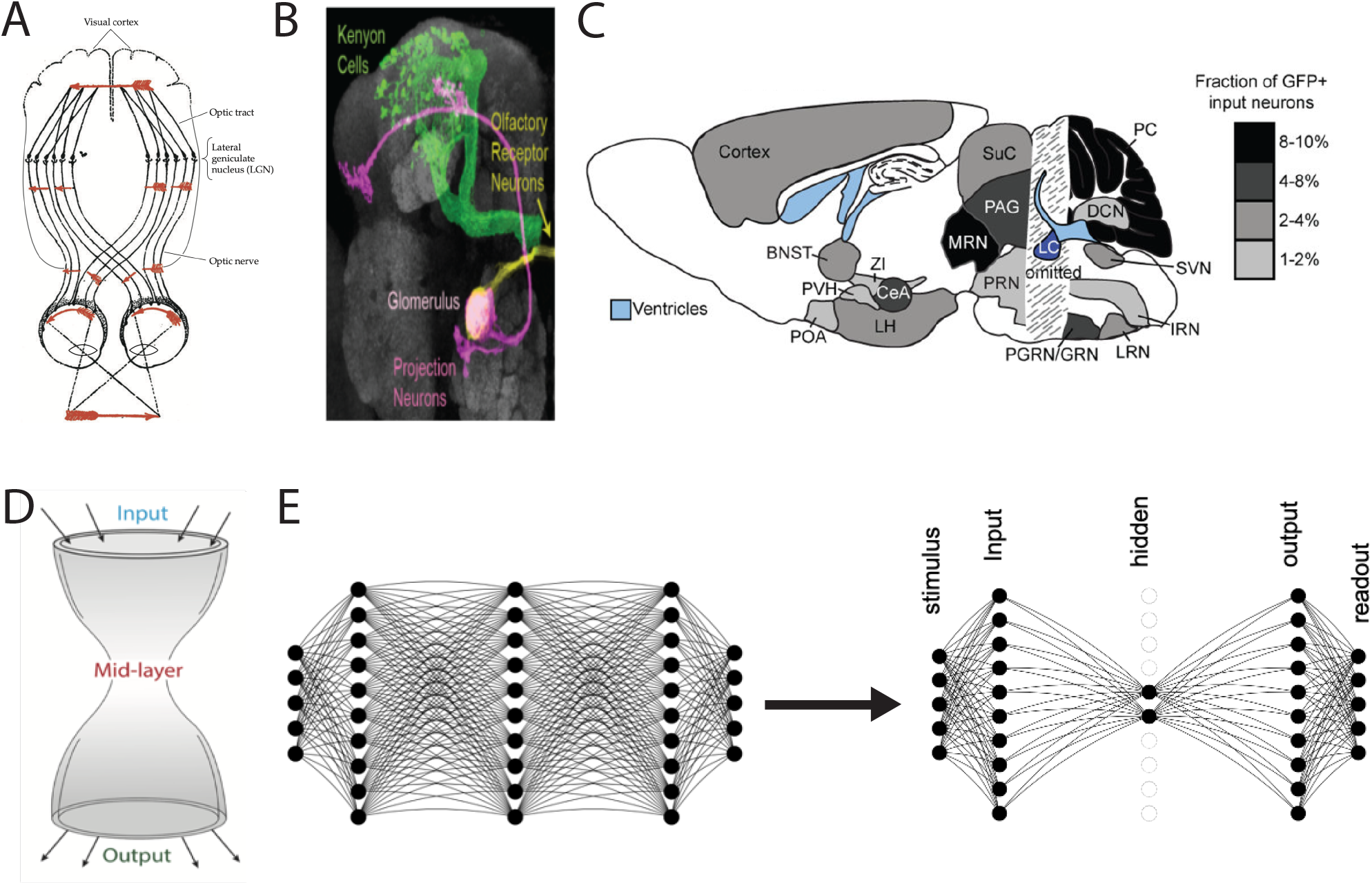
Observed and emergent bow-tie architecture (BTA) in neural networks. (A-C) BTA observed in visual (A), olfactory (B), and neuromodulatory (C) systems (with permission from (*17, 56, 57*)). (D) Schematic of BTA or ‘hourglass’ structure’ (*27*). (E) Transformation of a network from without BTA (left) to BTA (right) through training.

It has been shown that BTA, or part of such architecture, is useful for efficient information processing, including retaining more information (*23*), enhancing computational speed and accuracy (*24*), and amplifying variability of incoming stimuli while enhancing representation (*25*). In machine learning, BTA is widely used as an autoencoder for data feature compression or dimensional reduction, utilizing lower dimensional latent representation at the central bottleneck neural layer (*26*) (**Fig. 1D**, right). Despite the gradual uncovering of BTA functions in biological systems and applications in artificial neural systems, it is unclear how BTA can emerge.

Previous theoretical studies on BTA emergence were motivated by metabolic, gene regulatory or signalling systems instead of neural systems (*27, 28*). The networks considered were linear and network changes were due to variability or mutation of the network connections in simple multiplicative manner. Further, the models’ optimisation was simply to minimise the difference between the network’s matrix and some defined goal matrix. In comparison, neural networks in machine learning and neuroscience are generally nonlinear and connection weights or synaptic strengths are modified via learning rules and algorithms, and cognitive (e.g. decision-making) processing have to be readily and robustly performed (*29–31*). Critically, previous work trained neural networks in a predefined manner, which lacked sufficient generality.

In this work, we addressed the above issues by training feed-forward neural networks to perform various classification tasks on open standard datasets to understand how BTA can emerge in neural systems (**Fig. 1E**). We found non-negative network connections to be a key mechanism for BTA formation, and formally showed that this was achieved through amplification of error signals during learning and subsequent quenching of neural activities, especially at the hidden layer. Moreover, we found that the BTA became stabilized as the number of output neurons increased and the BTA wiring cost was low, hence, offering BTA’s functional advantages.

## Results

### Emergence of bow-tie architecture in neural network

To demonstrate the emergence of BTA, we first trained a five-layer neural network, with 500 neurons in each layer on a generic stimulus input masked by additive noise. Specifically, each stimulus or signal was represented by a 50-dimensional vector with numerical value *x* randomly sampled from a uniform distribution between 0 and 1, masked by a noise that followed an independent Gaussian noise *ϵ* with a mean of 0 and standard deviation of 0.1. The stimuli were pre-classified and banded into 100 classes based on the mean of each dimension. The network was trained to correctly classify a stimulus into the *i*^*th*^ class if the activity of the *i*^*th*^ readout neuron, described by a soft-max function, was greater than that of any other readout neuron given the same stimulus.

Initially, each neuron in a layer is fully connected to all neurons in the next layer, and the connection weight is randomly sampled from a uniform distribution *U* (0.01, 0.2). Each dimension of the stimulus vector acts as input to each neuron in the input (first) layer with predefined connection weights *W*_1_. The standard back-propagation learning algorithm (Materials and Methods) (*29*) was used to train the network to classify the stimuli into the predefined 100 class labels. We divided the dataset into two parts, 60% of which were for training and 40% for testing. The network was exposed to all stimuli during each epoch to classify the stimuli, with 256 samples per training batch. We purposefully kept the model training and testing procedures to be sufficiently simple to more clearly identify the underlying mechanism for BTA emergence.

**Fig. 2A-C** (left to right) shows snapshots of the evolution of the connection weights between the layers at the beginning, intermediate, and late stages during the training session. We can observe that as training progressed, the connection from stimulus-to-input layer *W*_1_, input-to-hidden layer *W*_2_, and hidden-to-output layer *W*_3_, became sparser while the strength of the remaining connections increased (**Fig. 2A-C**). After 20 training epochs, the classification accuracy for the training dataset was 0.9957, while it was 0.9955 for the testing dataset.

**Figure 2:**
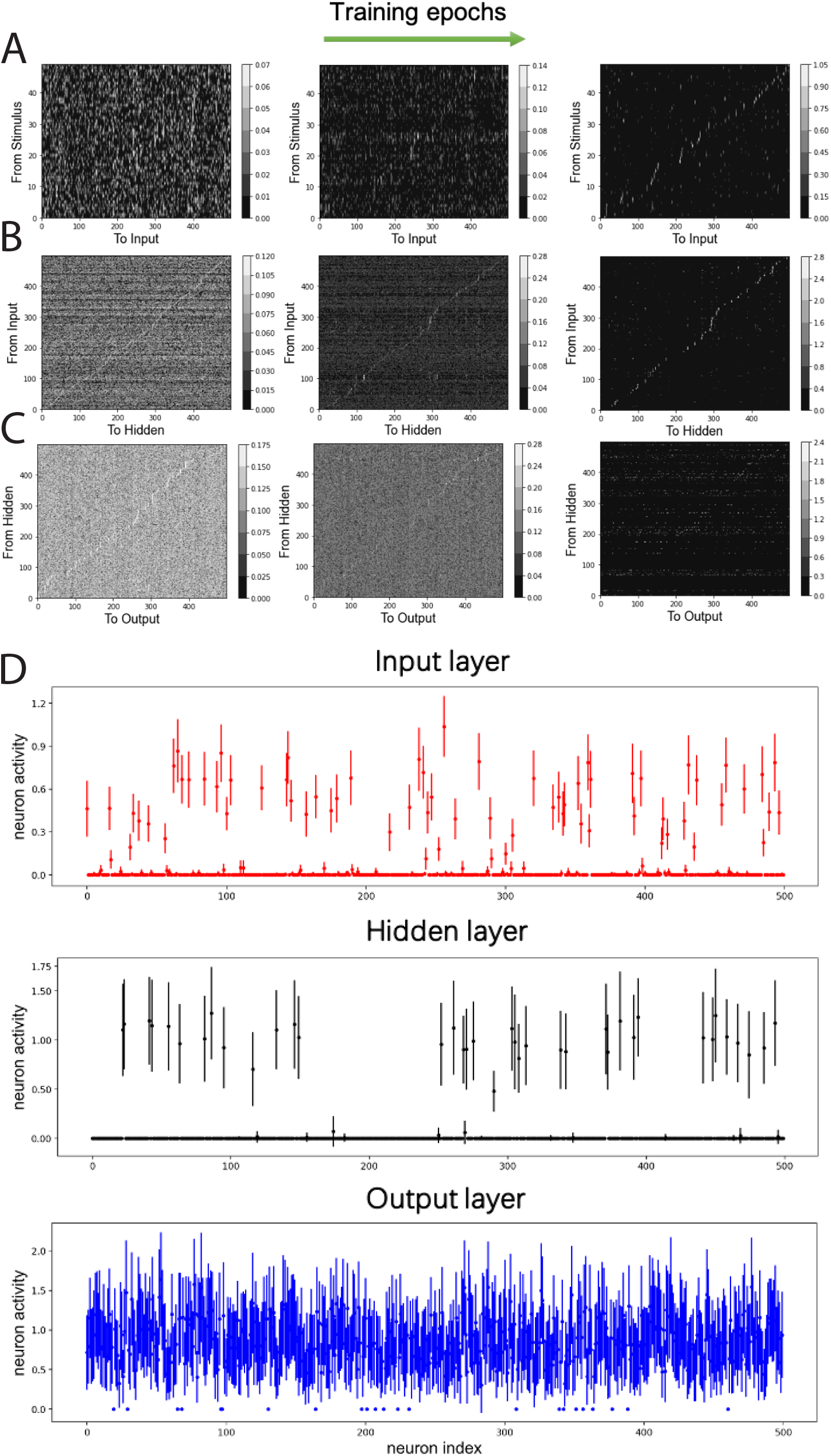
Neural network’s non-negative connection weights and neuronal activities due to training. (A-C) Connection weights from stimulus to input neurons (A), from input to hidden neurons (B), and from hidden neurons to output neurons (C), over training epochs (left to right). Color bars: values. (D) Neuronal activities of input, hidden and output neurons after training. Error bars: standard deviation across stimuli.

Next, the activity of the neurons in the input, hidden and output layers (*h*_1_, *h*_2_ and *h*_3_) are investigated. As higher neuronal activity variability across different stimuli can indicate the encoding of more stimulus information, we used the standard deviation of neuronal activity as an indicator for neuronal responsiveness (mean neuronal activity values gave similar results (not shown)). Specifically, by defining a neuron to be responsive or active if the standard deviation of its response to different samples was greater than 0.1, we found that the overall neuronal activities decreased after training, with the majority at the input and hidden layers being completely inactivated and not responding to the stimuli (**Fig. 2D**). More precisely, at the end of 50 training sessions with 20 epochs per session (in last training epoch), there were 225.69 ± 15.67 active input neurons, 153.31±20.79 active hidden neurons, and 498.85±1.29 active output neurons. Note also the relatively small variations in the numbers of active neurons across different training sessions.

Compared to all other layers, the hidden layer had the least number of active neurons after training (**Fig. 2D**, middle). This was also reflected in the lowest number of non-zero connection weights from the input layer to the hidden layer (**Fig. 2B**, right). Thus, we have shown the emergence of BTA in the network using a simple learning procedure on a generic classification task. Next, we shall formally demonstrate that the BTA occurs only with non-negative connections.

### Non-negative connectivity causes bow-tie architecture

The connection matrices from stimulus-to-input layer *W*_1_, input-to-hidden layer *W*_2_ and hiddento-output layer *W*_3_ were initially set as random matrices. Thus, the activities of the output neurons can be considered as random variables, since the different samples were transformed by a series of random matrices. However, the weighted sum of afferent inputs to a readout neuron (e.g., neuron c_4_ in **Fig. 3A**) from output neurons could initially just turned out to be larger than that of other readout neurons (e.g., neuron c_1_ in **Fig. 3A**), and hence the network would have a strong tendency to produce the same readout decision with different sample classes, i.e., poor classification accuracy (∼ 1%). As a consequence, the classification for all 256 samples in a batch in the first training epoch was concentrated to only one class (**Fig. 3B** right panel).

**Figure 3:**
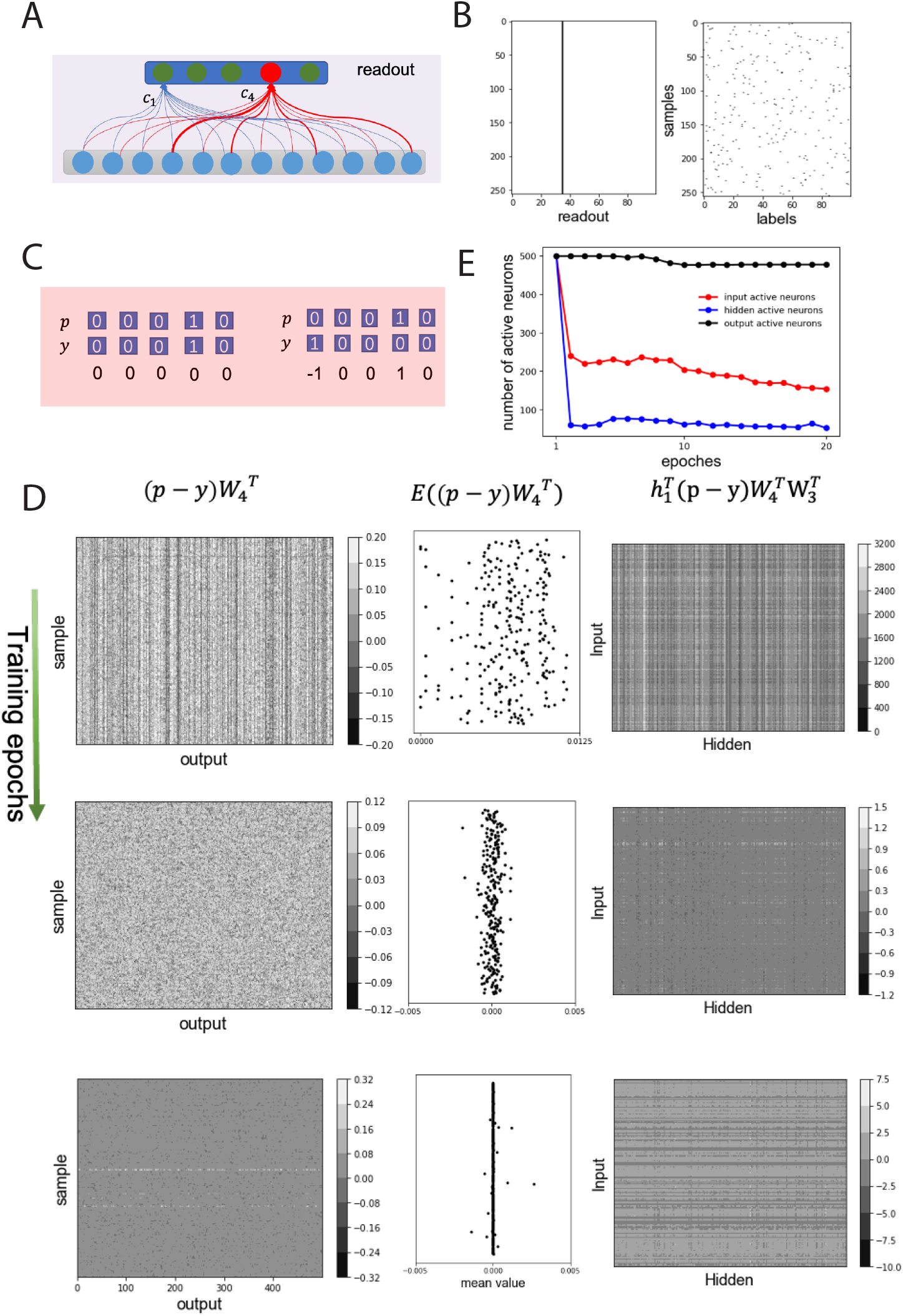
Non-negative connection weights amplify error signal and suppress neuronal activities, forming BTA. (A) Schematics of random initial connection weights from output to readout neurons. Weighted sum of afferents to readout neuron, c_4_ (red), larger than that of other neurons (e.g. neuron c_1_ (green)). (B) Class labels for each sample in one batch (left) and neuronal activity in read-out layer (right) in the first training epoch (same decision made regardless of samples). (C) Examples of classification decisions *p*, actual class labels *y*, and their errors *p* − *y*. Left (right): correct (error) decision. (D) Backpropagation of error signals. A random batch chosen and error signal of samples illustrated. Top, middle, bottom rows: first, middle, final training epochs. Left (right) columns: errors backpropagated from readout to output (from hidden to input) neurons; middle column: mean backpropagated error values from readout neurons. *W*_3_ (*W*_4_): connections from hidden to output (output to readout) layer. *h*_1_: input layer’s activity. (E) Number of active neurons in input (red), hidden (blue) and output (black) layers over training epochs.

Suppose for each sample, the readout layer of the neural network can be represented as a vector *p* (**Fig. 3C**, top), and the predefined class label of a sample is represented as a vector *y* (**Fig. 3C**, bottom) and randomly distributed in a batch (**Fig. 3B**, left). Assuming that the network accurately classifies the *i*^*th*^ sample, then the *i*^*th*^ neuron in the output layer will be activated, and the *i*^*th*^ element of decision vector *p* will be 1. This means that the decision vector *p* is the same as the predefined label of the sample *y*, i.e., *p* − *y* = 0. Then all elements of *p* − *y* are zero; thus, no error signal propagates back through the network layers, and the connection weights will not be changed. Otherwise, some elements of *p*−*y* are not zero. For example, **Fig. 3C** (right) shows the 1^*st*^ and 4^*th*^ elements of *p* − *y* to be −1 and 1, respectively, because the reported class of the sample is 4 (i.e., *p*_4_ = 1) but the predefined class of the sample is 1 (i.e., *y*_1_ = 1). This error signal will then be back-propagated to the output layer, the hidden layer, and the input layer. In particular, the error signal back-propagated to the output neurons depends on the efferent weights from the output neurons to the readout neurons, which should be in the form of a vector as 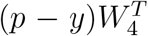, where 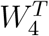 was the connectivity matrix for connections from output layer to readout layer (illustrated in **Fig. 3D**, left column) (*29*). Thus, the back-propagated error to the *m*^*th*^ output neuron is equal to the difference between the efferent weight from the *m*^*th*^ output neuron to the decision of the readout neuron (e.g., neuron *c*_4_ in **Fig. 3A**) and the efferent weight from the *m*^*th*^ output neuron to the readout neuron should be activated as in the predefined label.

Often, the total efferent weights from the output neurons to the activated readout neuron were greater than those to the inactive readout neurons, so the error signal induced by a probe should be non-negative. Considering 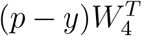 for each sample in the batch data as a random variable, the expectation value of 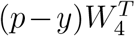 should be greater than zero for incorrect classification, or equal to zero for correct classification (**Fig. 3D**, middle column). According to the learning rule for connections from the hidden layer to the output layer, 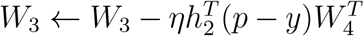 (Materials and methods, Eq. 2), with learning rate *η* and hidden layer (layer 3) activity *h*_2_, the total connection weight from the hidden layer to the output layer should decrease. Due to the non-negative connectivity constraint *W*_2_ and *W*_3_ for the earlier layers, the expectation of 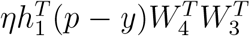 should be positive, and the error signal *p* − *y* being amplified by positive connections.

Further, the weights of the connections from the input to the hidden layer, *W*_2_, decreased according to the learning rule 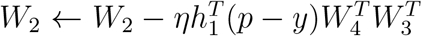. Considering that the stimulus is *s* ∈ [0, 1], the changed values of *W*_1_, i.e. 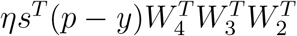, were usually smaller than that of *W*_2_. As a result, the connection weights from the input to the hidden layer, and from the hidden to the output layer decreased during the first training epoch.

Due to the decrease in the connection weights, the activity of the input and hidden neurons decreased significantly, and a portion of the input and hidden neurons became inactive in subsequent training epochs (**Fig. 3E**). As training progressed, the back-propagated error signals decreased (**Fig. 3D**, left and middle columns) and the magnitude of the weight change also decreased (**Fig. 3D**, right column), implying that the inactive neurons in the input and hidden layers remained unchanged (**Fig. 3E**; see also **Fig. 2D**). Considering that deafferentation can lead to the degeneration of the postsynaptic neurons in sensory systems (*32–34*), the inactive neurons in the system will degenerate and can be removed from the system. Therefore, the non-negative weights between layers significantly reduced the number of active neurons in the hidden layer, implying that the BTA had emerged from the network.

To further test the role of non-negative connectivity constraint on BTA formation, we trained a network with identical architecture and on the same classification task as above, but without the non-negative weight constraint. We first found that the trained network could perform the classification task as well, albeit with slightly better classification accuracy (0.9984 for the training dataset and 0.9964 for the testing dataset). We also found that when using a non-negative connectivity constraint, the network’s learning speed was slightly reduced. Particularly, if we defined the accuracy after the *n*^*th*^ epoch of training as *acc*(*n*), and the convergence rate *α* of training can be approximated as 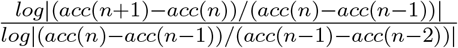 (*35*), we found *α* to be 0.850 and 0.825 for the network with and without non-negative connectivity constraints, respectively. Meanwhile, the trained connections between the layers were more dense without non-negative connectivity constraint (**Fig. 4A-C**), and neurons in the input, hidden and output layers were almost active (**Fig. 4D-F**; compared to **Fig. 2D**). Therefore, the training of the network without non-negative connectivity constraint did not result in the formation of BTA in the network.

**Figure 4:**
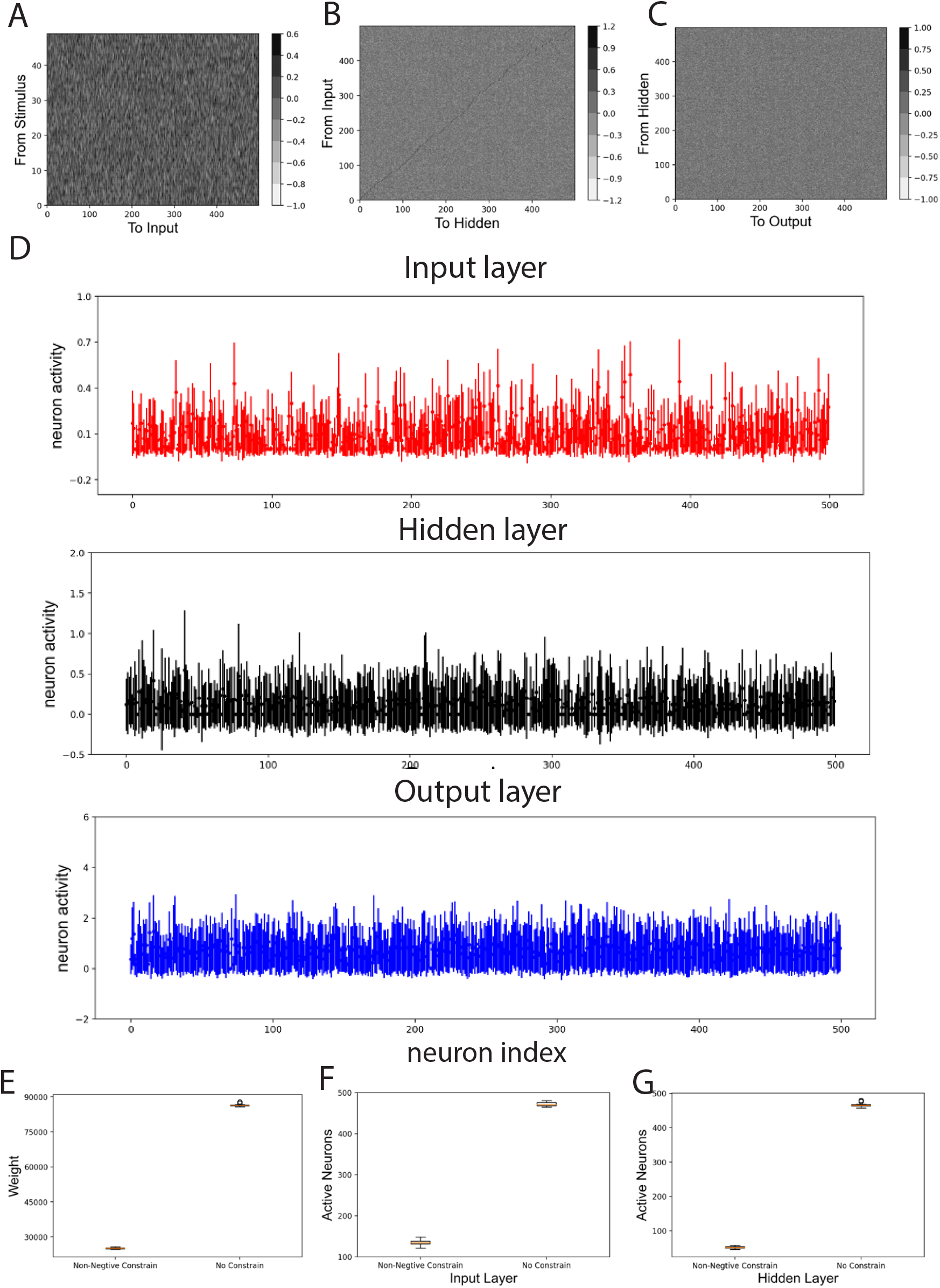
Sparse connectivity and low wiring cost of BTA. (A-C) Dense connection weights of network without non-negative connectivity constraint. (D-F) Activity of neurons without non-negative connectivity constraint. (G) Wiring cost of network with or without non-negative connectivity constraint. (H-I) Number of active neurons in input and hidden layers of network with or without non-negative connectivity constraint. Error bar: standard deviation over 10 differently initialized trained networks.

In summary, the non-negative connection weights between network layers amplified the error signal and suppressed the activities of a significant number of neurons in the input layer and especially in the hidden intermediate layer, resulting in the emergence of BTA. Moreover, the classification accuracy and training speed achieved by the non-negative connectivity constrained network, which resulted in BTA, was comparable to that of the network without such constraint.

### Bow-tie architecture is efficient, robust and generalizable

Since each synaptic connection is associated with metabolic cost (*36, 37*), we can evaluate the wiring cost for a neural network with BTA. For simplicity, we defined the total wiring cost of the connections as the sum of the absolute values of the network’s connection weights and then compared the wiring cost for a network with and without non-negative connectivity constraints. In **Fig. 4G-I**, it is clearly shown that the network with non-negative connectivity constraint had a much lower (about a third lower) connection cost than the one without this constraint. In addition, the number of active neurons in the input and the hidden layers of the network with BTA is much smaller than that in the network without BTA. These results suggest that networks with BTA may be highly energy efficient.

Although it may be desirable to have the BTA network to having a much smaller number of hidden neurons than other network layers, the hidden layer with its smaller structure may perhaps be more vulnerable to disruption or perturbation. To test the BTA’s robustness with respect to structural changes, we separately trained 10 networks with identical initial architecture but different initial connection weights to perform the same classification task. We then randomly removed active neurons in the input, hidden, and output layers of the trained network.

The removal of neurons generally degrades the classification accuracy. In particular, for the same proportion of active neurons removed in the input, the hidden, and the output layers, we found that removing the hidden neurons degraded the classification accuracy (**Fig. 5A**, black) more than that in the input or output layer (**Fig. 5A**, red and blue). This was expected because the number of active hidden neurons in the trained network was much smaller than the number of active input and output neurons. Moreover, the small number of hidden neurons may encode key latent features, and their removal can be costly to performance. Thus, active hidden neurons act as a critical network hub for processing important information, and their removal has greater effect than removing neurons from the other layers.

**Figure 5:**
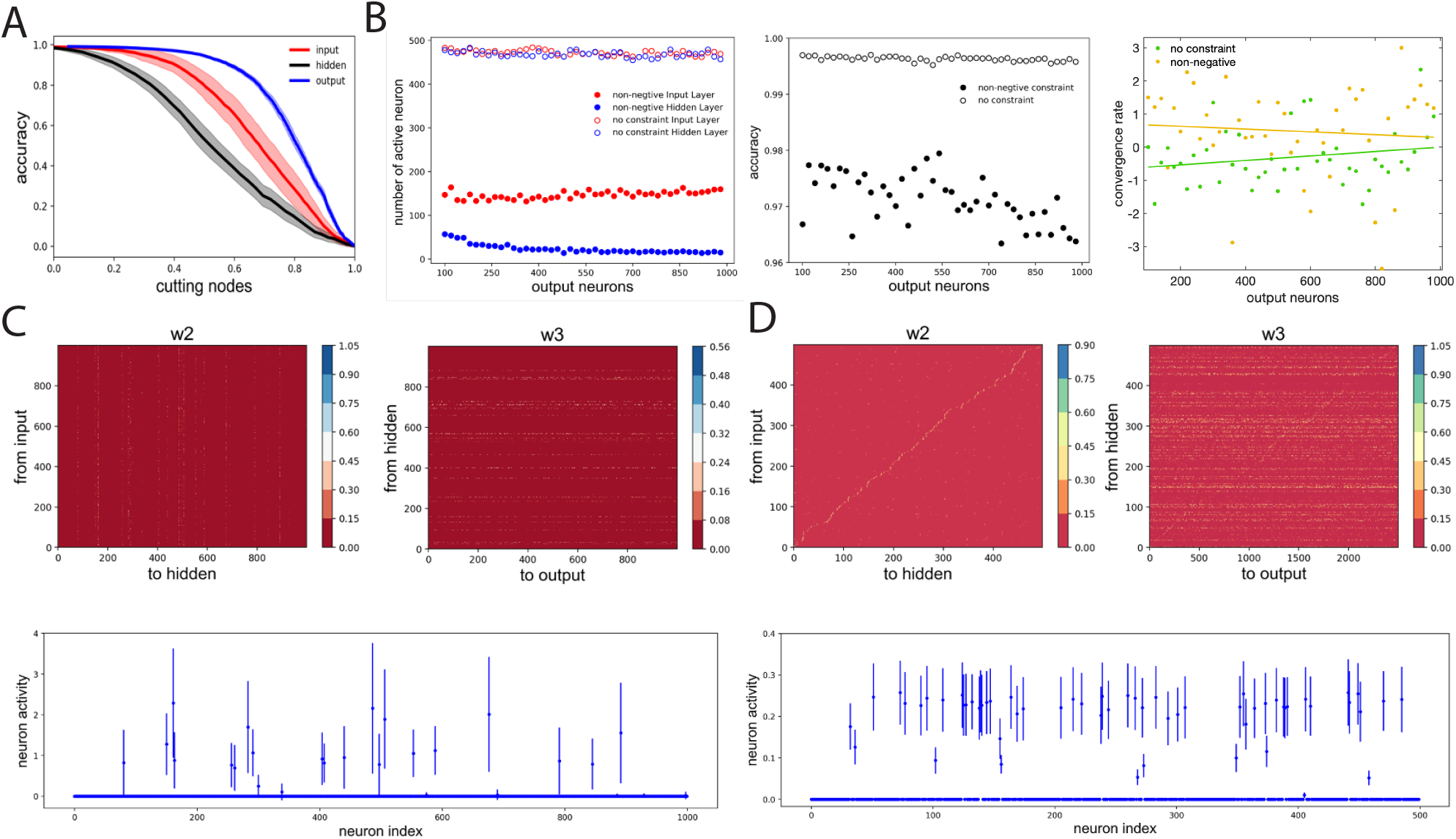
Robustness and generalizability of BTA. (A) Removal of neurons in the trained network at different layers decreased classification accuracy at different rates, with hidden neurons as the largest contributors. Horizontal axis: proportion of removed active neurons in input, hidden, or output layer. Shades: standard deviation of classification accuracy for 10 separately trained networks with different initial connection weights. (B) Left: with non-negative connectivity constraint, the number of output neurons did not substantially affect the final number of active input neurons (filled red circles) but decreased the active hidden neurons (filled blue circles). BTA was stabilised with increasing number of output neurons. Unfilled circles: without non-negative connectivity constraint. Middle, right: Classification accuracy (middle) and convergence rate (right) of networks with (yellow) and without (green) constraint. Fitted lines: linear regression *y* = 6.71 × 10^−4^*x* − 0.67 (green); *y* = −4.23 × 10^−4^*x* + 0.72 (yellow). (C) MNIST classification task. Trained connection weights from input to hidden layer (left) and from hidden to output layer (right) led to BTA. Small number of hidden neurons active after training (bottom). (D) Olfactory discrimination task. Trained connection weights from input to hidden layer (left), and from hidden to output layer (right) led to BTA. Small number of hidden neurons active after training (bottom).

As observed in **Fig. 3E** that the model’s active neurons in the output layer typically did not change relatively much, we next allowed the number of neurons in the output layer to vary from 100 to 1000, while fixing 500 input neurons and 500 hidden neurons in the network. The non-negative connectivity constraint was applied and each network structure was trained to perform the same classification task. We found that as the number of output neurons increased, the number of active input neurons did not vary too much, hovering around 140 (**Fig. 5B**, left, filled red circles), but the number of active hidden neurons was substantially reduced (**Fig. 5B**, left, filled blue circles). The BTA then became stabilized with further increase in the number of output neurons. Furthermore, increasing the number of output neurons did not have a substantial effect on the accuracy of classification accuracy (**Fig. 5B**, middle) and the convergence rate of the training process (**Fig. 5B**, right). Repeating the procedures on networks without the nonnegative connectivity constraint did not show substantial changes in the number of input and hidden neurons (**Fig. 5B**, left, opened filled circles). These results indicate that the BTA is robust to the variation in the number of output neurons, as had been observed in neural systems (*10, 38*).

So far, we have made use of a generic classification task using synthetically generated dataset. Hence, our next step was to investigate whether the emergence of BTA can be generalized to various more realistic classification tasks. Thus, we first trained a network with nonnegative connectivity constraint, initially with 1000 input neurons, 1000 hidden neurons, and 1000 output neurons to classify 10 handwritten digits in the well-known MNIST database with 60, 000 samples (*39*). We found the classification accuracy on the training and testing datasets to be high, at 0.9613 and 0.9260, respectively. We then repeated the training on the MNIST data for 50 networks, and found the numbers of active neurons were on average 380.14 ± 15.80, 87.92 ± 7.35, and 997.76 ± 1.3 for the input, hidden, and output layers, respectively. Hence, after training, the connections became more sparse (**Fig. 5C**) and the active neurons formed a BTA, similar to our observation with our synthetic generic dataset. Thus, the non-negative connectivity constrained network could classify very well with a robust emergent BTA.

Finally, we seek to know whether the BTA network can also perform odor discrimination task. Here, we trained the network with the same structure as in (*40*) but with a non-negative connectivity constraint to classify the 1, 008, 192 odor samples into 100 classes. This required a four-layer neural network, initially with 500, 500, 2500 and 100 neurons from the first odor stimulus layer to readouts (*40*). The accuracy of the training and testing datasets were found to be 0.9248 and 0.7865, respectively. After training, the network with non-negative connectivity constraint formed a BTA with sparse connectivity from input to hidden and from hidden to output neurons (**Fig. 5D**). Based on 50 sessions of training with 20 epochs per session, we found that the numbers of active neurons were 62.08 ± 3.71, 48.97 ± 4.19 and 2308.74 ± 39.60 for the input, hidden, and output layers of the network, respectively. This was again a BTA, consistent with previous work (*40*). Overall, we have shown that BTA emerged even with different, more realistic tasks, suggesting that BTA emergence could be generalized and applied to different cognitive tasks.

## Discussion

There is limited investigation on how BTA networks are formed, other than studies that have focused on metabolic, gene regulatory, or signaling systems (*41*). In contrast to the models used in these studies, neural networks are nonlinear, have both positive and negative connectivity, can perform cognitive tasks such as sensory discrimination, and can modify their architecture on a relatively faster timescale through learning and synaptic plasticity (*42*). How BTA generally arises in neural networks and the conditions that allow it to do so have not been systematically addressed and mechanistically explained.

In this work, we identified a cause of BTA formation in the training of neural networks in the context of noisy signal classification and discrimination tasks. Specifically, we found that training neural networks with non-negative connections in the neural layers led to BTA. This was achieved by amplifying back-propagated error signals during network learning while silencing neuronal activity within the input layer, and to a greater extent, the hidden layer (**Fig. 1**). This finding is supported by evidence that there are more excitatory than inhibitory neurons in the cortex, possibly for optimal computation (*43*), and there are also more long-range excitatory (positive) than inhibitory (negative) synapses (*44, 45*).

Our results were tested on various discrimination tasks and class sizes; from 10 classes for handwriting discrimination to 100 classes for the generic noisy signal and odor discrimination tasks. We demonstrated that BTA formation can occur in neural networks regardless of discrimination class size or tasks. In addition, we showed that BTA required less wiring than non-BTA networks without sacrificing much in terms of discrimination accuracy and learning speed. With fewer active neurons and sparser connectivity, BTA networks with non-negative connectivity constraints require fewer wiring resources than without the constraint (**Fig. 4**), and thus can attain lower metabolic costs and more energy-efficient information processing.

Given that neural systems are generally heterogeneous and changing, it is not clear whether and how BTA neural network functions can be affected by structural variability. Here, we showed that discrimination accuracy was robust, with only a gradual reduction from the ablation of neurons in any input or output layer in the trained network, with the hidden layer exerting the strongest functional influence (**Fig. 5A**). Hence, bottleneck-like brain nuclei such as the ventral tegmental area with dopamine-producing neurons, the raphe nuclei with serotonin-producing neurons, and thalamic nuclei may be more vulnerable and susceptible to perturbation and disruption, and subsequently, affect brain functions or states, or even lead to brain disorders (*46–49*).

The present study is limited in terms of using a specific (error back-propagating) learning algorithm, simple neural networks with feedforward architecture, and sensory discrimination tasks. These simplified assumptions allow theoretical tractability and explainability on how non-negative connectivity can lead to BTA. Future work will extend the present work’s limitations by investigating whether the concept and principles of the BTA neural network formation found in this study can be also be obtained with other neural network architectures, different learning rules and cognitive functions.

To summarize, we have shown non-negative connections in neural networks to be a key cause of BTA emergence. We have also shown that neural networks with BTA have efficient wiring costs, are relatively robust to network size, and could perform well in different discrimination tasks.

## Materials and methods

### Data description

Three datasets were used for the classification tasks. For the first dataset, we synthetically generated a total of 2 million samples. Each sample *s* = (*s*_1_, *s*_2_, …, *s*_50_) is composed of signal *x* and noise *ϵ*, i.e. *s*_*i*_ = *x*_*i*_ + *ϵ*_*i*_, where *x*_*i*_ ∼ *U* (0, 1) and *ϵ*_*i*_ ∼ *N* (0, 0.05). For each stimulus *s*_*i*_, we generated 20, 000 samples, and we divided them into 100 categories or classes based on the mean value of the samples.

To verify the robustness of the BTA’s performance, we used two other open datasets. One of them is the well-established MNIST (*50*) with 60, 000 samples of handwritten digits. The other is the odor detection data (*40*), where each odor is denoted as *s* = (*s*_1_, *s*_2_, …, *s*_50_) and *s*_*i*_ ∼ *U* (0, 1) with a total of 1, 008, 192 samples, and the samples were categorized into 100 classes according to their nearest neighbors (*51*).

### Neural network model and architecture

Feedforward neural network model architecture was used in this study. The neurons were distributed in 5 layers, namely, they are the stimulus, input, hidden, output, and readout layers (**Fig. 1e**, left). The input layer of neurons represents sensory-based neurons, such as the retina in the visual system or neurons with olfactory receptors in the olfactory system. The hidden layer of neurons can be considered as an intermediate processing stage, similar to the ganglion cells in the visual system or projection neurons in the olfactory system. The output layer of neurons represents the primary visual cortical neurons or Kenyon cells in the visual or odor systems, respectively. The readout layer of neurons encodes the classes, with the number of readout neurons equaling the number of classes to be discriminated or identified by the model.

The activity of each layer was represented by a vector as *h*_0_, *h*_1_, *h*_2_, *h*_3_ and *h*_4_ for the respective 5 layers in the network. Each activity vector had a dimension 1 × *n*_*i*_ where *n*_*i*_ indicated the number of neurons in the *i*^*th*^ layer. We let matrix *W*_1_ describe the connections from the stimulus layer to the input layer, and *W*_2_, *W*_3_ and *W*_4_ for connections from the input to the hidden, from the hidden to the output, and the output to the readout layer, respectively. For simplification, we set *h*_0_ = *s* where *s* is the stimulus. We used the nonlinear rectified linear unit (ReLU) function to describe the activity of the input, the hidden, the output, and the readout neurons, mathematically described by 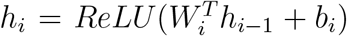, where *i* = 1, 2, 3, 4, and *b*_*i*_ is a bias vector.

### Model training and testing

The network was initialized as a fully connected network and the initial connection weights followed a uniform distribution between 0.01 and 0.2. We used a standard backpropagation (BP) algorithm to train the network to classify the stimuli into different classes, with support regarding its approximation in neurobiological systems (*52*). We chose cross entropy as the loss function (*53*): *Loss* = − Σ_*j*_ *y*_*j*_*log*(*p*_*j*_), with 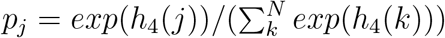, where *y* is the predefined label of stimulus in vector form, *p*_*j*_ the soft-max function of the *j*^*th*^ readout neuron. The connection weights for consecutive iterations were updated as follows:

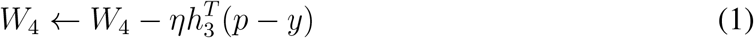

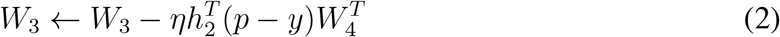

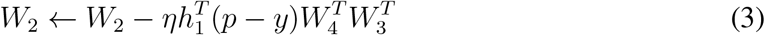

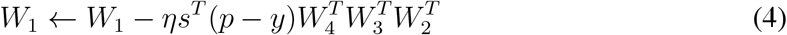

where *η* = 0.1 is the learning rate. We followed the stochastic gradient descent method (*54*) and used the Keras Adam optimizer to train the network (*55*). In the presence of non-negative constraint, *W*_1_, *W*_2_ and *W*_3_ were set to be non-negative constrain,namely, with *W* = 0 if *W <* 0.

### Model parameters and performance on three tasks

Table 1 summarizes the model parameters, training, and other information such as the number of classes, input dimension, batch size, number of training epochs, number of neurons, convergent rate, and discrimination (F-1) accuracy.

**Table 1:** Training details for three tasks with non-negative constraint. The data for the training results is based on the average of 50 training sessions.

|  | Synthetic dataset | MNIST dataset | Odor detecting dataset |
| --- | --- | --- | --- |
| No. classes | 100 | 10 | 100 |
| Input dimension | 50 | 784 | 50 |
| Batch size | 256 | 512 | 256 |
| No. epochs | 20 | 200 | 20 |
| Training set size | 600000 | 36000 | 1000000 |
| Testing set size | 400000 | 24000 | 8192 |
| Initial neurons | 500, 500, 500 | 1000, 1000, 1000 | 500, 500, 2500 |
| Convergence rate | 0.850 | 0.307 | -0.318 |
| Accuracy of training set | 99.57% | 96.13% | 92.48% |
| Accuracy of testing set | 99.55% | 92.60% | 78.65% |
| No. active neurons | 225.69, 153.31, 498.85 | 380.14, 87.92, 997.76 | 62.08, 48.97, 2308.74 |
| F-1 score | 0.9699 | 0.9707 | 0.7436 |

## Data analysis

We recorded the activity of each neuron exposed to different samples and calculated the mean and variance of the activity over the samples. To investigate the effects of non-negative connectivity constraints, we trained 10 networks (training more networks yielded similar results) with non-negative connectivity constraint given different initialized connection weights to perform the same classification task. We then trained another 10 network without a non-negative connectivity constraint to perform the same classification task. After training, we compared the connectivity of these networks and the activity of neurons.

To analyze the robustness of the BTA, we trained 10 networks with different initialized connection weights to perform the same classification task using the synthetic, generic noisy signal dataset. After training, we removed some active input neurons, hidden neurons, and output neurons from the trained network. We had the ablated network perform the same classification task and obtained the accuracy of the classification.

The source codes, generated data, and analyses that support the findings of this study will be made available upon publication.

## Acknowledgements

This work was funded by National Science Foundation of China (NSFC) under grant 32171094 (D-HW). KW-L was supported by HSC R&D (STL/5540/19) and MRC (MC OC 20020).

